# POT1a depletion in the developing brain leads to p53-dependent neuronal cell death and ataxia

**DOI:** 10.1101/335059

**Authors:** Robert She, Charlie Clapp, Eros Lazzerini Denchi

## Abstract

The processes that control genome stability are essential for the development of the Central Nervous System (CNS) and the prevention of neurological disease. Here we investigated whether activation of a DNA damage response by dysfunctional telomeres affects neurogenesis. More specifically, we analyzed ATR-dependent DNA damage response by depletion of POT1a in neural progenitors. These experiments revealed that POT1a inactivation leads to a partially penetrant lethality, and surviving mice displayed ataxia due to loss of neuronal progenitor cells, and died by 3 weeks of age. Inactivation of p53 was sufficient to completely suppress the lethality associated with POT1a depletion and rescued the neuronal defects characterizing POT1a depleted animals. In contrast, activation of ATM by the depletion of the shelterin protein TRF2 resulted in a fully penetrant lethality that could not be rescued by p53 inactivation (Lobanova et al.). This data reveals that activation of distinct types of DNA damage response pathway give rise to different types of neuropathology. Moreover, our data provides an explanation for the heterogeneity of the neurological defects observed in patients affected by telomere biological disorders.

## Introduction

Mutations affecting DNA damage response factors are associated with neurodegenerative disease in humans. For example, inactivation of the DNA damage kinase ATM (Ataxia telangiectasia mutated) is associated with ataxia (for review see (Shiloh and Ziv, 2013)). Similarly, mutations in the ATR (ATM and Rad3-related) kinase are the cause of Seckel syndrome, a disease characterized by microcephaly and mental retardation (O’Driscoll et al., 2003). ATM and ATR are activated in response to DNA lesions and orchestrate cell cycle arrest, activation of DNA repair, and apoptosis. These kinases are activated by different type of lesions with ATR induced by single-stranded DNA while ATM is activated by DNA double-strand breaks (DSBs) (Cimprich and Cortez, 2008; Shiloh and Ziv, 2013).

Telomeres are essential nucleoprotein structures that protect chromosome ends from being recognized as sites of DNA lesions. In mammals, suppression of the DNA damage response at chromosome ends is achieved by the recruitment of a specialized protein complex termed shelterin that binds the telomeric DNA repeats (TTAGGGn) (for review, see (de Lange, 2009)). Telomeric repeats are lost upon cellular division leading to progressive telomeric shortening with age. When telomeres become critically short, they fail to recruit sufficient shelterin complex. Dysfunctional telomeres are recognized as sites of DNA damage leading to aberrant DNA repair and activation of the tumor protein p53 which in turn depending on the cellular context can trigger apoptosis or senescence (for review see (Denchi, 2009)). The current model for the mechanism of POT1 mediated suppression of ATR signaling involves the ability of POT1 to bind the single-stranded portion of telomeric DNA (Denchi and de Lange, 2007a; Gong and de Lange, 2010). When POT1 is depleted, the single stranded binding protein RPA localizes to the telomeric g-overhang leading to ATR activation. In contrast, the model for TRF2-mediated suppression of ATM involves the formation of a protective secondary structure called T-loop that sequester the ends of a chromosome, thereby preventing its recognition as a double-stranded break (DSB) (Doksani et al., 2013; Griffith et al., 1999). In addition, TRF2 is able to actively suppress localization of the DNA damage factor RNF168 thus suppressing the activation of a DNA damage response downstream of ATM activation (Okamoto et al., 2013).

In humans, there are a number of syndromes characterized by short telomeres that collectively are referred to as telomere biology disorders (TBDs) (for review see (Savage, 2014)) which include Dyskeratosis congenita (DC), Hoyeraal–Hreidarsson syndrome and Coats plus syndrome. These complex diseases range in clinical severity, from DC, which involves multiple organs, to diseases with only one organ affected, such as pulmonary fibrosis. In these patients, the most affected tissues tend to be those with high turnover, like the hematopoietic system. This is consistent with the notion that telomere erosion occurs more rapidly in these tissues. However, defects in organs such as the brain, which is not characterized by a high turnover rate, are also observed in these patients. For instance, patients affected the Hoyeraal–Hreidarsson syndrome show severe cerebellar hypoplasia and microcephaly (Hoyeraal et al., 1970). Moreover, a significant fraction of Dyskeratosis congenita patients have cognitive defects (Rackley et al., 2012). These observations suggest that telomere dysfunction could affect brain development or proper neurological function. Additionally, late generation telomerase knockout mice display defects in tissues with a high rate of turnover as well as neurological impairment and cognitive defects (Ferron et al., 2004). Furthermore, telomerase reactivation in mice with short telomeres results in telomere elongation and the alleviation of neurological impairments (Jaskelioff et al., 2011; Tomas-Loba et al., 2008). Thus, it is anticipated that upon critical telomere shortening, the initiation of a DNA damage response at chromosome ends would lead to either cell death or impairment in neurological function. This was confirmed by the observation that depletion of the shelterin component TRF2 in the developing mouse brain leads to severe perturbation in neurogenesis and causes cognitive defects (Lobanova et al.). Notably, TRF2 depletion in nestin-positive progenitor cells causes the death of the vast majority of neuronal cells and results in embryonic lethality (Lobanova et al.). In contrast, telomere dysfunction induced by depletion of the shelterin component POT1a has a mild effect on brain development and does not result in embryonic lethality (Lee et al., 2014). Moreover, POT1a depletion is well tolerated in the vast majority of neuronal cells and only affects a subset of neurons in the cerebellum (Lee et al., 2014). It is currently unclear why there is such a difference in the phenotypes induced by depletion of TRF2 and POT1a. A major difference between the telomere dysfunction induced by TRF2 depletion and POT1a depletion is the level of end-to-end chromosome fusions induced.

Notably, two distinct conditional knockout mice for POT1a have been generated, one in which the first coding exon is depleted following Cre expression (Wu et al., 2006). In this setting, an alternative start site generates a truncated protein which is predicted to localize at telomeres (for more details see (Hockemeyer et al., 2006)). This mouse was used to probe the consequences of POT1a depletion during neuronal development (Wu et al., 2006). A different POT1a conditional knockout mouse was generated by the de Lange laboratory in which the third coding exon can be depleted generating truncation products that cannot localize to telomeres (Hockemeyer et al., 2006). We have previously used this conditional knockout mouse to test the role of POT1a depletion in the lymphoid compartment (Pinzaru et al., 2016). Here, we investigated whether depletion of POT1a using this mouse model results in similar phenotypes as the ones previously reported (Lee et al., 2014). Our data confirm that conditional depletion of POT1a in the developing mouse brain affects primarily the cerebellum, a phenotype that can be fully rescued by p53 inactivation. Strikingly however, our results indicate that POT1a depletion using the POT1a allele generated by the de Lange laboratory (Hockemeyer et al., 2006) has a more penetrant and severe phenotype compared to what reported using the other available POT1a conditional knock out mouse model (Wu et al., 2006). Collectively, this data reveals that different levels of telomere dysfunction can give rise to profoundly different outcomes in the developing brain.

## Material and methods

### Mice

The *Pot1a* and *p53* conditional alleles have been previously described (Celli and de Lange, 2005; Hockemeyer et al., 2006; Marino et al., 2000). To achieve inactivation of the POT1a and p53 in the nervous system Nestin-Cre mice (Marino et al., 2000) (B6.Cg-Tg(Nes-Cre)1Kln/J, JAX#003771) were used to generate: Pot1a^Flox/Flox^;Nestin-cre (referred as *Pot1a*^*F/F,Nes*^) and Pot1aFlox/Flox; p53^Flox/Flox^; Nestin-cre (referred as *Pot1a*^*F/F*^*p53*^*F/F,Nes,*^). After establishing the different mouse lines, the genotyping was performed by Transnetyx. All animals were maintained in accordance with the National Institutes of Health’s Guide for the care and use of laboratory animals. The institutional animal care and use committee at The Scripps Research Institute approved all procedures for animal use.

### Histology

Histology procedures were performed as previously described (Denchi et al., 2006) (Lobanova et al.). Brains were collected at indicated time points and fixed by perfusion with 4% buffered PFA. All samples were paraffin-embedded and sectioned by a microtome (4μm). Sections were stained with Gill modified hematoxylin (Harleco) and eosin Y (Ricca Chemicals) to detect morphological differences. For IHC, paraffin-embedded sections were treated for 20 minutes in boiling EDTA buffer for antigen retrieval. DNA damage response activation and apoptosis detection was performed by the antibodies anti-phospho-H2AX (mouse, 1:3000, Millipore) and anti-Casp3 (rabbit, 1:1000, Cell Signaling). An anti-doublecortin antibody (rabbit, 1:3000, abcam) was used to label neuronal progenitors. Detection was performed using the ImmPRESS HRP Polymer Detection Kit (Vector Labs) and the Metal Enhanced DAB Substrate Kit (Thermo Scientific) according to the manufacturer’s instructions. Each staining was performed on at least three samples of each genotype. All slides were examined and imaged using an Axio Imager M1 microscope (Zeiss).

## Results

### POT1a depletion in neural progenitor cells leads to ataxia

The mouse genome encodes for two POT1 orthologs: POT1a and POT1b (Hockemeyer et al., 2006; Wu et al., 2006). POT1a is involved in protection of chromosome ends from the DNA damage response pathway, while POT1b regulates the amount of single-stranded DNA at the telomere terminus (Hockemeyer et al., 2006). To define the consequences of telomere deprotection in the developing central nervous system (CNS) we crossed conditional *Pot1a* knock out mice (*Pot1a*^*F/F*^) with a Nestin-Cre (Nes) line, which expresses Cre in neural progenitors. The resulting *Pot1a*^*F/F,Nes*^ mice were born in sub-Mendelian ratio. The vast majority (approx. 90%) of the expected *Pot1a*^*F/F,Nes*^ mice died before birth. The fraction of *Pot1a*^*F/F,Nes*^ mice that successfully completed embryogenesis appeared normal at birth (Fig 1a-b). However, by two weeks of age, all *Pot1a*^*F/F,Nes*^ animals were significantly smaller in size and showed evidence of severe ataxia (Fig 1c and Supplemental video 1). All the *Pot1a*^*F/F,Nes*^ mice had to be euthanized by 21 days of age due to moribund state. The brains of the two week old *Pot1a*^*F/F,Nes*^ animals were significantly smaller than those of the control mice (Fig 1d) and histological examination revealed a striking reduction in the development of the cerebellum and of the dentate gyrus (Fig 1d). The severe cerebellar hypoplasia observed in *Pot1a*^*F/F,Nes*^ mice explains the lack of motor skills in these animals and is reminiscent of other mouse models with increased levels of DNA damage in the developing brain, such as ATR and TOPBP1 conditional knockout mice (Lee et al., 2009; Lee et al., 2012). These results are in line with the induction of a DNA damage response following POT1a depletion (Denchi and de Lange, 2007b; Gong and de Lange, 2010; Guo et al., 2007) and suggests that during embryogenesis the cerebellum is particularly susceptible to DNA damage induction.

**Figure 1.**
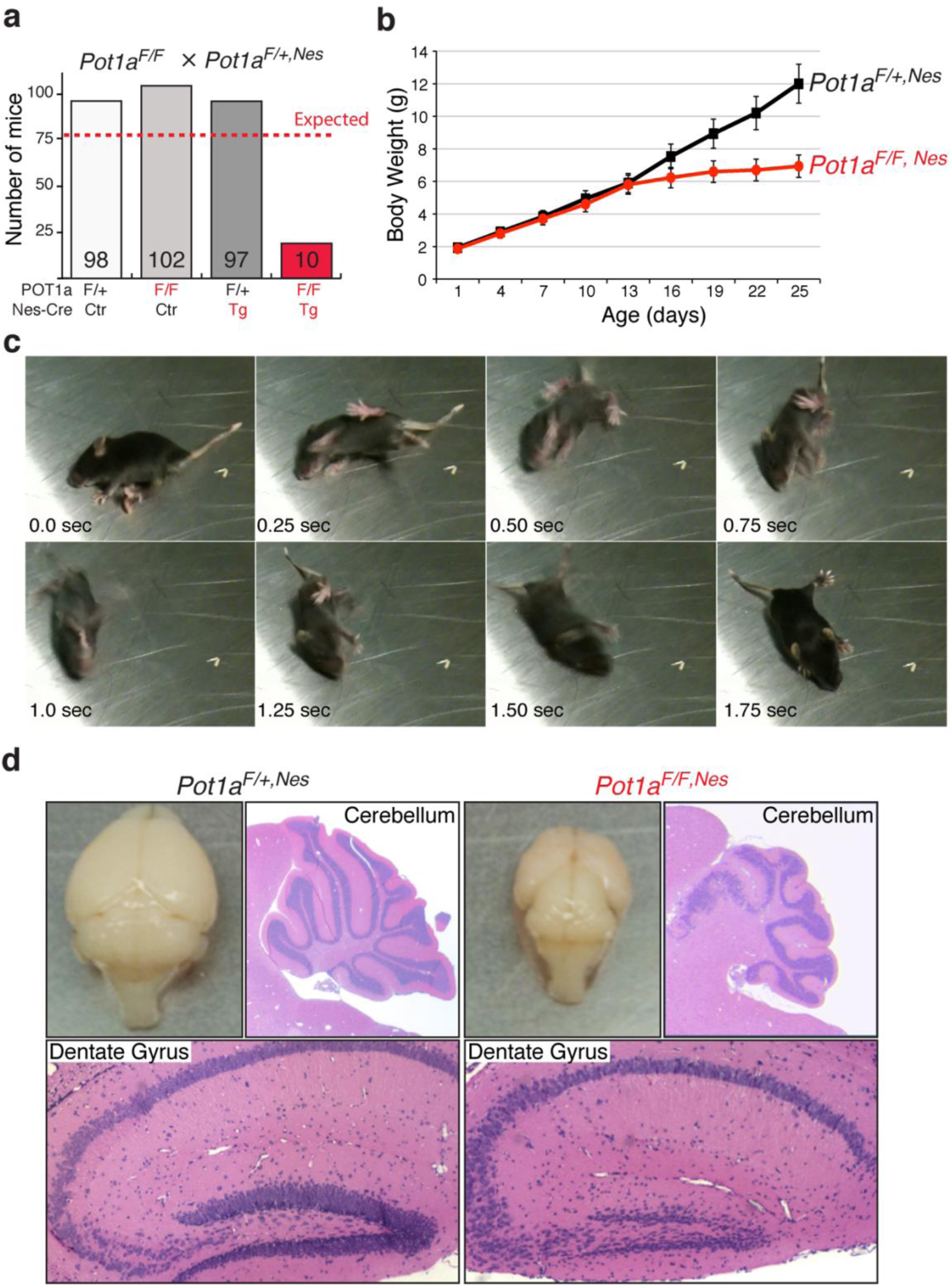
*POT1a is required for* in some cases, as a driver for genomic *development.* **a,** Birth and expected survival rates of offspring of all genotypes from selected breeding couple *Pot1a*^*F/F*^ X *Pot1a*^*F/+,Nes*^. **b,** Body weight curves of control (*Pot1a*^*F/+,Nes*^) mice (n=3) and of *Pot1a*^*F/F,Nes*^ mice (n-3). **c,** Still frames of a *Pot1a*^*F/F,Nes*^ displaying ataxia phenotype at 17 days of age. **d,** Picture of the brain of the mouse shown in panel c and of a littermate control mouse at day 17 of age. Adjacent to it are H&E stainings of *Pot1a*^*F/F,Nes*^ mice and control animals showing hypoplasia of the cerebellum and impaired formation of the dentate gyrus.

### Phenotypes associated with Pot1a depletion in neural progenitor can be rescued by p53 inactivation

The DNA damage response pathway results in p53 activation and, in turn, apoptosis is induced. Therefore, we decided to test whether the lethality and neuronal degeneration in *Pot1a*^*F/F,Nes*^ mice was caused by p53 activation. To this end, we crossed *Pot1a*^*F/F,Nes*^ mice with mice carrying a conditional *p53* allele (Marino et al., 2000). The resulting *Pot1a*^*F/F*^*p53*^*F/F,Nes*^ mice were born at the expected mendelian ratio, were viable, and appeared morphologically indistinguishable from control mice (Fig 2a-c). We followed a cohort of *Pot1a*^*F/F*^*p53*^*F/F,Nes*^ mice and relative control mice for 10 months and observed no evidence of increased malignancy associated with concomitant inactivation of POT1a and p53 in the CNS (Fig 2a). In contrast, depletion of POT1a in animals heterozygous for p53 (*Pot1a*^*F/F*^*p53*^*F/+,Nes*^) leads to a phenotype that was indistinguishable from the depletion of POT1a from p53 wild type animals (Fig 2a-c). Next, we assessed whether p53 inactivation suppressed the developmental defects observed in the *Pot1a* ^*F/F,Nes*^ mice. We found that *Pot1a*^*F/F*^*p53*^*F/F,Nes*^ mice had the same size brain as POT1a proficient mice (Fig 3a). In addition, p53 inactivation was sufficient to restore the proper morphology and size of the cerebellum (Fig 3b) and of the dentate gyrus (Fig 3c). In accordance with this, *Pot1a*^*F/F*^*p53*^*F/F,Nes*^ mice displayed no signs of ataxia (Fig 2b). Moreover, p53 inactivation suppressed the loss of DCX-positive progenitor cells that is characteristic of *Pot1a* ^*F/F,Nes*^ mice (Fig 3c). This data shows that upon POT1a depletion, p53 induction—rather than telomere dysfunction — causes embryonic lethality and neuronal degeneration. Next, we tested whether the lethality associated with TRF2 depletion in nestin-positive cells (Lobanova et al.) was caused by p53 induction. Our data shows that p53 inactivation was not sufficient to rescue the lethality associated with TRF2 inactivation (Fig 2e). This data further highlights the difference between consequences of POT1 and TRF2 depletion in neural progenitor cells.

**Figure 2.**
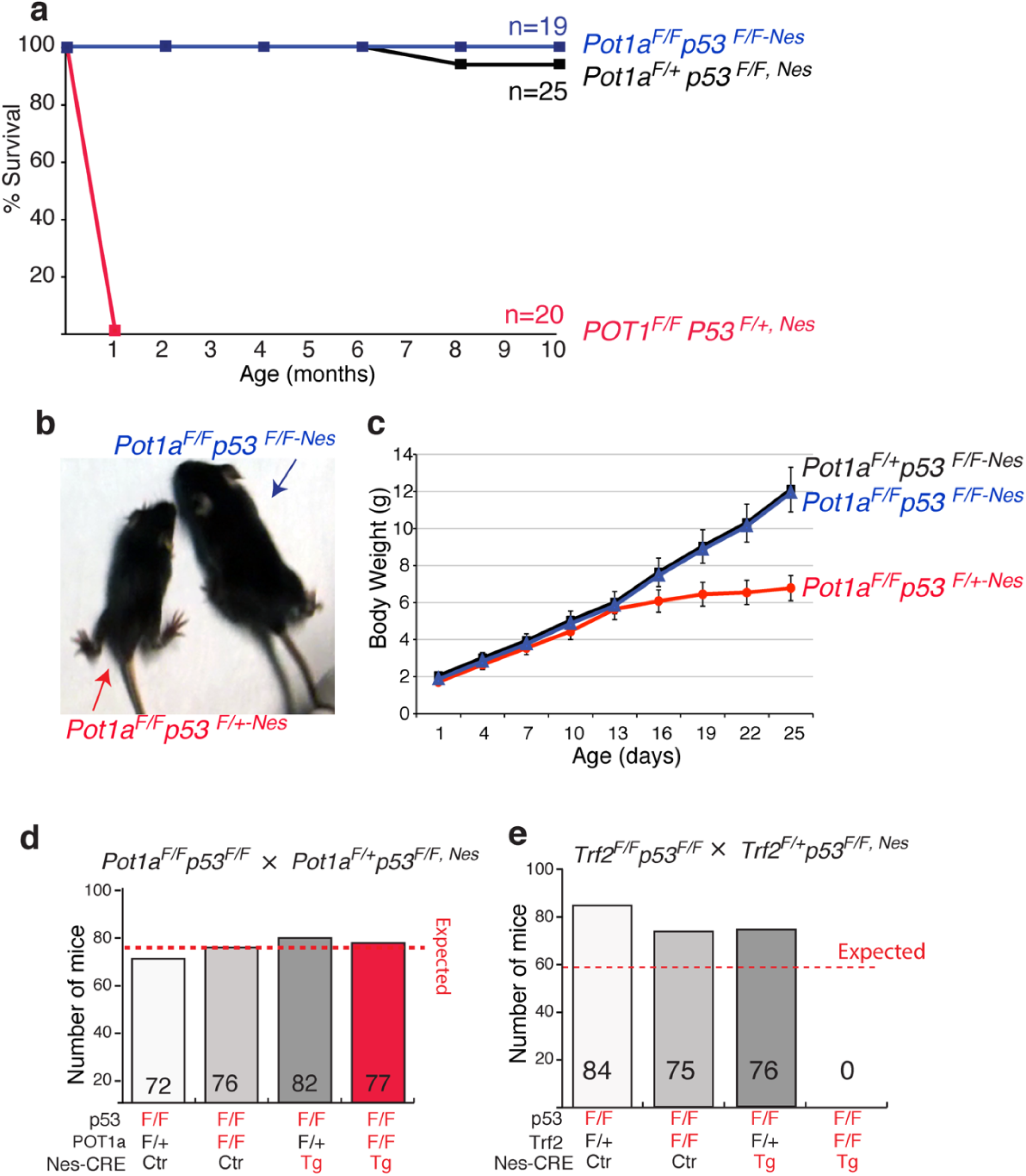
p53 *depletion suppresses lethality associated with POT1a depletion in the CNS.* **a,** Survival curve of control (*Pot1a*^*F/+*^p53^*F/F,Nes*^) mice (n=25), of POT1 depleted (*Pot1a*^*F/F*^p53^*F/+,Nes*^) mice (n=20) and of double knockout (*Pot1a*^*F/F*^p53^*F/F,Nes*^) animals (n=19). **b,** picture of a POT1 depleted (*Pot1a*^*F/F*^p53^*F/+,Nes*^) mouse and of a littermate double knockout (*Pot1a*^*F/F*^p53^*F/F,Nes*^) mouse. **c,** Body weight curves of female control (*Pot1a*^*F/+*^p53^*F/F,Nes*^) mice (n=3), of POT1 depleted (*Pot1a*^*F/F*^p53^*F/+,Nes*^) mice (n=3) and of double knockout (*Pot1a*^*F/F*^p53^*F/F,Nes*^) (n=3). *Lethality* caused by *TRF2 loss in the developing CNS cannot be rescued by p53-inactivation.* **d,** Birth and expected survival rates of offspring of all genotypes from selected breeding couple (*Pot1a*^*F/F*^*p53*^*F/F*^ X *Pot1a*^*F/+*^*p53*^*FF, Nes*^). **e,** Birth and expected survival rates of offspring of all genotypes from selected breeding couple (*Trf2*^*F/F*^*p53*^*F/F*^ X *Trf2*^*F/+*^*p53*^*FF, Nes*^).

**Figure 3.**
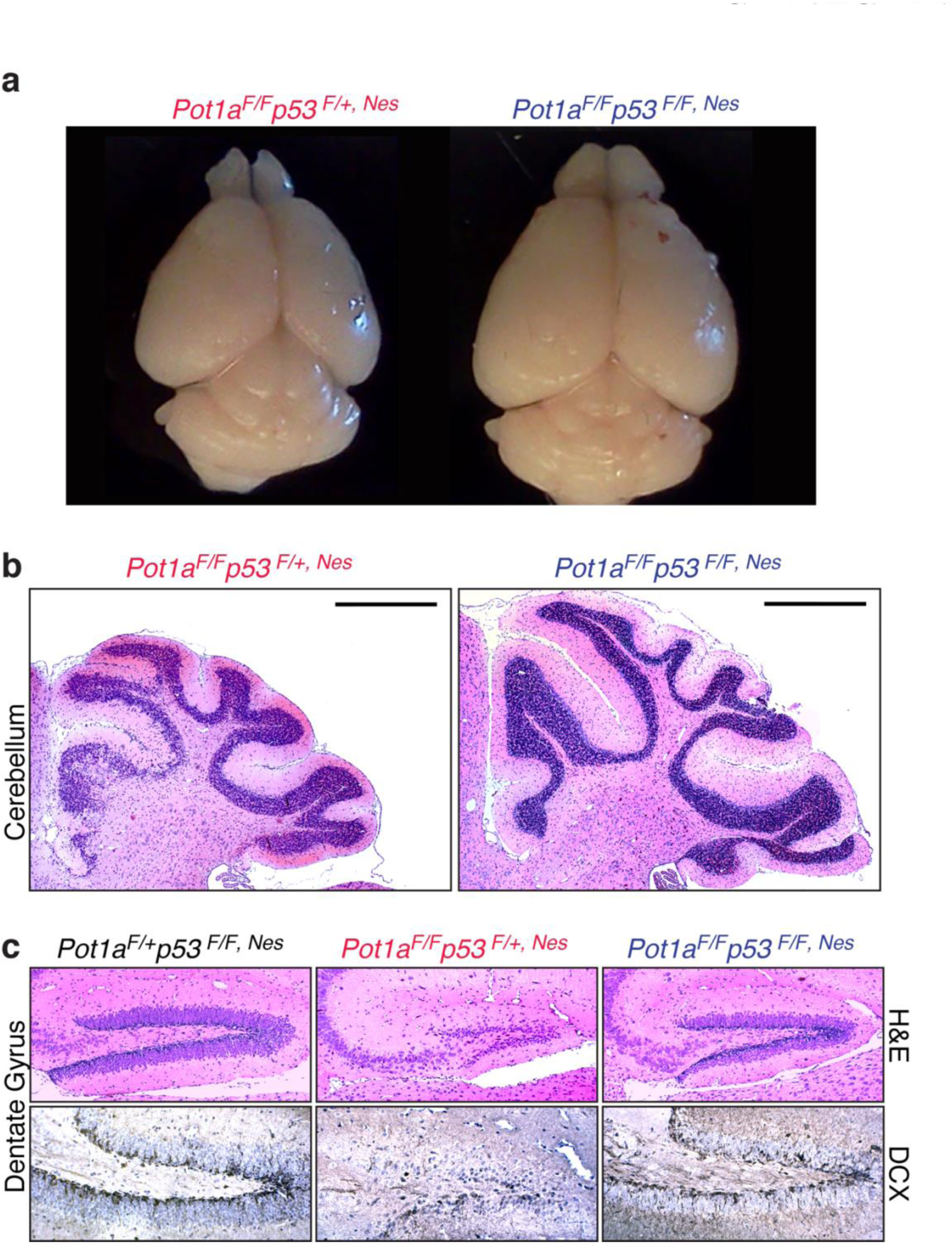
p53 *depletion suppresses neuronal loss associated with POT1a depletion in the CNS.* **a,** Representative pictures of brains isolated from 17 day old, *Pot1a*^*F/F*^p53^*F/+,Nes*^and *Pot1a*^*F/F*^p53^*F/F,Nes*^ littermate controls. **b,** H&E staining showing cerebellular hypoplasia in *Pot1a*^*F/F*^p53^*F/+,Nes*^ mice and its suppression in *Pot1a*^*F/F*^p53^*F/F,Nes*^ animals. Scale bars represent 1 mm. **c,** H&E staining and Immunohistochemistry (IHC) for doublecortin (DCX), a marker of newly born neurons in the dentate gyrus. Note the reduction in the granule neurons and in DCX-positive cells in *Pot1a*^*F/F*^p53^*F/+,Nes*^ mice compared to double knockout (*Pot1a*^*F/F*^p53^*F/F,Nes*^) mice.

## Discussion

### Acute telomere dysfunction affects neurogenesis

In the context of highly proliferative tissues, telomere dysfunction arising from critical shortening of telomere repeats has been established has a major factor limiting regeneration potential and, in some cases, as a driver for genomic instability. Less is known on the potential role of telomere dysfunction in organs that do not undergo high levels of turnover. Some lines of evidence suggested that critically short telomeres may cause neurological defects in human pathologies, but the incomplete penetrance of neurological phenotypes in telomere biology disorders (TBDs) raised the possibility that telomere dysfunction by itself may not be sufficient to cause these phenotypes. Mouse models lacking telomerase have been well characterized and demonstrated that animals with critically short telomeres have neurological associated defects, such as decreased sense of smell and cognitive defects (Rackley et al., 2012). However, in these mouse models, telomere dysfunction is not restricted to the CNS, thus there is the possibility that systemic factors might contribute to these phenotypes. Recently, conditional depletion of shelterin factors has been used to define the consequences of telomere dysfunction in the CNS (Lee et al., 2014) (Lobanova et al.). In these models, deletion of POT1a or TRF2 are restricted to the CNS using the neuronal specific Cre lines Nestin-Cre; Nex-CRE and Dcx-CRE. The results of these studies revealed a striking difference in the phenotype triggered by POT1a depletion and TRF2 depletion. Indeed, while depletion of POT1a using the Nestin-CRE line resulted in limited neurodegeneration, TRF2 depletion in the same cell types produce massive neurodegeneration and resulted in embryonic lethality. Here, we used a different mouse strain than the one previously reported to conditionally deplete POT1a in nestin positive cells. Our results revealed the POT1a depletion profoundly impacts brain morphology and neurogenesis. Only a small fraction (approx. 10%) of animals completed development and reached birth. Further, animals that reached the neonatal stage developed ataxia and died by the age of three weeks. This data highlights the importance of proper telomere protection for neurogenesis and neuronal function. The phenotypes observed in our study are more severe than the ones previously reported by the McKinnon laboratory but importantly share the same fundamental properties. In both cases, the only affected area is the cerebellum, and in both cases, p53 inactivation fully rescues the observed degeneration. Observed differences are unlikely to be caused by differences in genetic background since in both studies mice were in a Bl6 background. Additionally, both studies used the same CRE driver line, excluding possible differences in the efficiency of CRE-mediated depletion. The major contributing difference is related to the conditional POT1a knockout mouse strain used. The McKinnon laboratory used a conditional knockout mouse model in which the first coding exon of POT1a is depleted following Cre expression (Wu et al., 2006). In this setting, an N-terminally truncated POT1a protein lacking about one third of the first OB-fold can be generated using an alternative ATG on the second coding exon (see for complete detail the supplemental information in (Hockemeyer et al., 2006)). This protein is predicted to localize at telomeres through the interaction with the telomere associated protein TPP1. In contrast, we used a conditional KO mouse in which, upon CRE expression, the third coding exon is depleted, leading to the formation of a small C-terminally truncated POT1a allele that cannot localize to telomeres and it is therefore fully not functional (Hockemeyer et al., 2006).

### p53 –dependent and –independent neuronal loss upon telomere dysfunction

A striking finding in our study was the effect of p53 inactivation on the consequences of telomere dysfunction. Firstly, we found that p53 inactivation could fully rescue the phenotype of POT1a depletion in the CNS. This finding shows that the lethality associated with POT1a depletion in neuronal progenitors is caused by activation of a DNA damage response that triggers p53 activation. Strikingly, this data shows that once the p53 response is abrogated, telomere dysfunction initiated by POT1a depletion can be well tolerated in the CNS. TRF2-depletion, which has been shown to display a more severe phenotype, could not be rescued by p53 inactivation. There are two main differences between POT1a and TRF2 depletion that could explain the differential p53-dependent response: i) TRF2 depletion leads to frequent end-to-end chromosome fusions (Celli and de Lange, 2005; Denchi and de Lange, 2007a; Doksani et al., 2013; Karlseder et al., 1999; Lazzerini Denchi et al., 2006) that could affect the division or differentiation of neuronal progenitor cells. In contrast POT1a depletion leads only to modest levels of telomere fusions (Denchi and de lange, 2007c; Gong and de Lange, 2010; Hockemeyer et al., 2006; Hockemeyer et al., 2005; Wu et al., 2006), which could be well tolerated in the CNS. ii) TRF2 depletion triggers an ATM-dependent DNA damage response while POT1a depletion triggers an ATR-dependent DNA damage response (Denchi and de Lange, 2007a). It is therefore possible that activation of these kinases might lead to differences in phenotypes arising from telomere dysfunction. However, this seems unlikely given the established interconnection between these two signaling pathway. Indeed, work by the McKinnon lab established that in the CNS, POT1a depletion leads to ATR activation that sequentially triggers ATM activation. Based on this data, we favor the hypothesis that the level of telomere fusions determines the outcome of telomere dysfunction in the developing brain.

### Telomere dysfunction and neurological defects

Our data indicates that the level of telomere dysfunction dictates the degree of neuronal loss in the developing brain. The microcephaly and cerebellar hypoplasia observed in *Pot1a*^*F/F,Nes*^ mice resemble the phenotypes observed in patients affected with the Hoyeraal**–**Hreidarsson syndrome (Hoyeraal et al., 1970). This data suggests that these patients suffer from telomere dysfunction occurring in neuronal progenitors, likely causing the observed brain development defects. In contrast, patients with other telomere biology disorders such as DC patients show either no neurological defect or less severe phenotypes (Rackley et al., 2012). It is conceivable that neuronal progenitors of DC patients have sufficiently long telomeres to maintain telomere protection and ensure a brain development. Nevertheless, some DC patients do display some cognitive defects (Rackley et al., 2012), suggesting that telomere dysfunction arising in more differentiated cell types or during adult neurogenesis could affect the proper neuronal function. Future experiments employing conditional TRF2 and POT1a depletion in differentiated neurons will be required to test the role of telomere dysfunction in these cell types.

Finally, the finding that microcephaly and cerebellar hypoplasia of *POT1a*^*F/F,Nes*^ can be fully rescued by p53 inactivation suggest that some of the pathologies of patients affected with the severe Hoyeraal**–**Hreidarsson syndrome could be suppressed. Future experiments will be required to fully explore this possibility and to provide potential avenues of treatment. One key factor would be to establish whether transient suppression of the p53 pathway may relieve some of the developmental defects associated with this disease. This would be extremely important given the central role for p53 for tumor suppression. To this end further experiments will be required to test whether this transient suppression of p53 during development might be sufficient to bypass the lethality associated with POT1a loss during brain development.

## Acknowledgement

We would like to thank Titia de Lange for proving the *Pot1a* conditional knockout mice and Uli Muller for providing the Nestin-Cre line. We are grateful to the Lazzerini Denchi lab helpful discussion. This work was supported by grants from the NIH (AG038677) and the American Cancer Society (RSG-14-186) to E.L.D. and from the Julia Brown Research Scholarship for Health and Medical Professions (R.S.).

